# Analysis of nucleus and cytoplasm-specific RNA fractions demonstrates that a significant proportion of the genetic regulation of gene expression across the human brain occurs post-transcriptionally

**DOI:** 10.1101/2022.06.28.497921

**Authors:** Karishma D’Sa, Sebastian Guelfi, Jana Vandrovcova, Regina H. Reynolds, David Zhang, John Hardy, Juan A. Botía, Michael E. Weale, Sarah A. Gagliano Taliun, Kerrin S. Small, Mina Ryten

## Abstract

Gaining insight into the genetic regulation of gene expression in human brain is key to the interpretation of genome-wide association studies for major neurological and neuropsychiatric diseases. Expression quantitative trait loci (eQTL) analyses have largely been used to achieve this, providing valuable insights into the genetic regulation of steady-state RNA in human brain, but not distinguishing between molecular processes regulating transcription and stability. RNA quantification within cellular fractions can disentangle these processes in cell types and tissues which are challenging to model in vitro. We investigated the underlying molecular processes driving the genetic regulation of gene expression specific to a cellular fraction using allele-specific expression (ASE). Applying ASE analysis to genomic and transcriptomic data from paired nuclear and cytoplasmic fractions of anterior prefrontal cortex, cerebellar cortex and putamen tissues from 4 post-mortem neuropathologically-confirmed control human brains, we demonstrate that a significant proportion of genetic regulation of gene expression occurs post-transcriptionally in the cytoplasm, with genes undergoing this form of regulation more likely to be synaptic. These findings have implications for understanding the structure of gene expression regulation in human brain, and importantly the interpretation of rapidly growing single-nucleus brain RNA-sequencing and eQTL datasets, where cytoplasm-specific regulatory events could be missed.

## INTRODUCTION

Over the last 10 years genome-wide association studies (GWAS) have successfully identified risk loci for the major neurological and neuropsychiatric diseases (1-5). However, in common with most complex diseases, the risk loci identified have been located in non-coding genomic regions. Since these loci are generally thought to affect disease risk through changes in gene expression, understanding the genetic regulation of gene expression in human brain has become an area of key interest. To date, this challenge has been primarily met through expression quantitative trait loci (eQTL) analyses (6-12), which have correlated genotype to gene expression in order to identify loci with regulatory potential. However, as these studies measure total gene expression levels in post-mortem human brain, they cannot differentiate between the molecular processes driving changes in mRNA abundance, namely the effect of a variant on transcriptional rate as compared to another process, such as degradation.

In this context, it is important to recognise that mRNA abundance is the result of multiple dynamically regulated events taking place across different cellular compartments. Briefly, the life cycle of an mRNA molecule starts in the nucleus where DNA is transcribed to form pre-mRNA. This is followed by the post-transcriptional processes of 5’-end capping, splicing, 3’-end cleavage and polyadenylation to generate mature mRNA, which is exported into the cytoplasm through nuclear pores. In contrast, mRNA molecules that are incompletely processed in the nucleus are retained, as our many long noncoding RNAs (lncRNAs) (13,14). While the former undergo degradation (15,16), the lncRNAs remain stable (13,14). Once in the cytoplasm, the mRNA either undergoes translation into protein within the cell body or may be transported to a specified subcellular location for local translation. In neurons, which are often large and structurally complex cells in the brain, RNA localization is increasingly being recognised as a key post-transcriptional process with a role in synaptic plasticity (17-20). Finally, mRNA molecules undergo degradation through a range of processes (15,21) including mRNA decay, which involves shortening of the poly(A) followed by decapping and degradation (16), and the nonsense mediated decay pathway (commonly triggered by premature stop codons). Collectively, these processes from transcription to degradation are key to maintaining mRNA abundance and must be tightly controlled for cell survival.

While a number of foundational eQTL analyses have led to the assumption that the majority of the regulatory sites operate by changing transcriptional rate (22), there is evidence to support the role of genetic regulation at other stages of the RNA life cycle (23,24). This includes genetic regulation of alternative splicing (25-28), polyadenylation (29,30), mRNA stability, translation, localization by 3’UTRs (31,32) and degradation (33). However, many of these analyses require the use of in vitro model systems which prevent their application to human brain and so the molecular processes driving eQTLs in brain specifically have largely remained unknown. While the use of iPSC-derived neurons could enable relevant time course experiments to be performed, it is well documented that iPSC-derived neuronal cultures and even organoid cultures differ significantly from human brain, in terms of the maturity they achieve and their cellular complexity, making it difficult to validate their relevance. An alternative is to use post-mortem brain tissue directly, studying the regulation of RNA processes occurring in the natural cellular compartments (namely, the nucleus and cytoplasm) to broadly assess the landscape of genetic regulation of gene expression.

Allele-specific expression (ASE) analysis is a means of assessing the genetic regulation of gene expression (10,34) which is well-suited to this type of experimental approach and has already been used for this purpose in cell lines (13). ASE analysis measures differential mRNA expression levels of 2 alleles of a heterozygous variant and so is a within individual analysis which is not affected by common confounding factors. Rather the power to infer an ASE at a heterozygous site is largely dependent on the read depth at that site (35-37). Importantly, pairing the nuclear-cytoplasmic RNA fractionation process within a sample ensures that biases affecting one cellular fraction would also operate on the other, so increasing the power of this analysis (13). In this study, we performed ASE analysis on nuclear and cytoplasmic RNA fractions from 3 brain regions (anterior prefrontal cortex, putamen and cerebellar cortex) derived from 4 individuals. These regions were selected as they are anatomically distinct and important to human disease (10,38-45). Together this unique sample set was used to investigate the relative importance of nucleus- and cytoplasm-specific processes in the genetic regulation of gene expression across the human brain.

## MATERIAL AND METHODS

### Generation and processing of RNA sequencing data

Post-mortem samples dissected from the anterior prefrontal cortex, putamen, and cerebellar cortex and originating from 4 donors were obtained from the MRC Sudden Death Brain and Tissue Bank. The brain tissue samples were collected from neuropathologically normal individuals of European descent, ranging in age from 41-57 (Supplementary Table 1). All samples had fully informed consent for retrieval and were authorized for ethically approved scientific investigation (Research Ethics Committee number 10/H0716/3).

Paired nuclear and cytoplasmic total RNA was extracted from each tissue sample using the Norgen Cytoplasmic and Nuclear RNA purification kit (Norgen Biotec Comp, Canada) according to the manufacturer’s specifications. Following DNase I treatment, the separate cellular fractions were obtained by centrifugation and RNA extracted using spin column chromatography. Cytoplasmic RNA samples were assessed for RNA integrity using Agilent 2100 Bioanalyzer. Nuclear and cytoplasmic RNA was used separately as input for Illumina’s TruSeq stranded total RNA library preparation kit with Ribo-Zero (Illumina, USA), and the resulting cDNA libraries underwent 100bp paired end sequencing (Illumina HiSeq4000). Real time base call and quality checks were performed using Illumina HiSeq Real Time Analysis software and fastq files were generated with Illumina’s CASAVA software to obtain ∼234M reads per sample.

Fastp (v 0.20.0), a fast all-in-one FASTQ pre-processor, was used for adapter trimming, read filtering and base correction (46) (v0.20.0) with parameters set allowing automatic detection of adapters, correction of bases in case of mismatches in overlapping regions of paired-end reads, exclusion of reads with lengths shorter than 36bp and checks for sequence over-representation. Default settings were used for the remaining parameters. Processed reads were mapped to the GRCh38 human reference genome via STAR (v 2.7.0a) using gene annotations from Ensembl v97 (47). An average of ∼234M reads per sample were processed (∼193M reads in the nuclear and ∼275M in the cytoplasmic samples) with an average of 96.2% reads per sample passing quality filters.

Parameters were set to match ENCODE options, as described in the STAR manual. Accordingly, the maximum number of mismatches per pair relative to the read length was set as 0.04. Unlike ENCODE, only reads that mapped to a single location in the genome were included. Post-alignment quality metrics were generated using RSeQC (48) (v2.6.4) and MultiQC (49) (v1.8.dev0). We found that an average of 87.45% of reads were uniquely mapped. There was a higher percentage of reads mapping to intronic regions in nuclear samples compared to the corresponding cytoplasmic samples. This was expected due to the higher levels of pre-mRNA in the nuclear fraction. Intronic reads were also observed in the cytoplasmic fractions and this could have been due to both the presence of unannotated transcripts and some RNA contamination from the corresponding nuclear fractions. Pipeline source code can be found in https://github.com/RHReynolds/RNAseqProcessing.

### DNA extraction and genotyping

Using 100-200mg of cerebellar tissue from each individual, DNA extraction was performed using Qiagen’s DNeasy Blood & Tissue Kit (Qiagen, UK) according the manufacturer’s protocol. This was followed by the generation of whole exome sequencing libraries using Nextera Rapid Exomes 12 and sequencing on the Illumina HiSeq4000 to generate an average of 80.3 million reads per sample. The resulting data was aligned to the human genome build hg19 with Novoalign and variant calling was performed using GATK (version 3.2-2) haplotype caller. Individual level gvcfs were merged into batches of 200 samples (covering 7,513 Exomes in total), and genotyping across all batches was then performed with the GATK joint genotyping tool. Variant sites were filtered using VQSR, with variants in the 99.9% tranche and above included (high quality variants). Genotypes were marked as missing if the reported genotype had a GQ<=15 or there were <4 reads (or >1000 reads) covering the site. Variant quality checks were performed and variants with quality of depth < 2 were excluded. The resulting vcf had 35,954 bi-allelic sites for the 4 individuals. DNA originating from each of the 4 donors was also analysed using the Omni Express Exome-8 Kit. Data from these individuals were batched together with that from 134 individuals that had been previously processed (6,50). The SNPs were assessed for strand ambiguity (http://www.well.ox.ac.uk/~wrayner/tools/#Checking) and those with evidence of ambiguity, as well as indels were removed. Finally, SNPs with MAF>5%, HWE of 0.0001 were retained, resulting in 745,784 loci. The WES and genotype data for each individual were merged giving preference to variant calls originating from WES data. The final vcf file containing 853,243 SNPs were uploaded to the Sanger Imputation Service (51) and phased with Eagle2 (52) using the Haplotype Reference Consortium (release 1.1) (51) as the reference panel. The resulting phased vcf contained an average of 268,240 heterozygous SNPs (hetSNPs) per individual after lift over of the coordinates from hg19 to hg38 using the UCSC online tool (53).

### Generation of gene level expression measures and differential gene expression analysis

Transcripts were quantified in each RNA-Seq sample using Salmon (54) (v0.14.1). We used the mapping-based mode in which the raw reads were aligned to the reference transcriptome, Ensembl version 97. Salmon was run using its default variational Bayesian Expectation-Maximisation algorithm with parameters to correct for sequence, non-uniform coverage biases (like 5’ or 3’ bias) and GC bias in the data. The bootstrapping option was set to enable Salmon to assess the technical variation in the abundance estimated. The R package tximport (55) was used to convert transcript-specific expression values measured in transcripts per kilobase million (TPM) to gene-level quantification values. Following gene-level quantification, the principal component analysis (PCA) analysis was performed using DeSeq’s plotPCA() function to visualize sample differences. Investigating the sources of variability (56) we found that tissue (region) correlated best with the 1st and 3rd axes, fractionation (condition) correlated with the 2nd axis and RIN with the 4^th^ (Supplementary figure 1). Differential gene expression across cellular fractions was conducted using DeSeq2 (57) controlling for individual effects (∼individual ID + fraction). Genes were considered significantly differentially expressed if FDR < 0.05. The apeglm method (58) was applied in line with DeSeq2’s recommendation to generate accurate log_2_FoldChange estimates. The Enhanced volcano (59) R package was used to generate the volcano plots, for genes with FDR < 5% and > 2 fold expression.

### Assessment of fractionation quality

Contamination between the nuclear and cytoplasmic fractions was investigated using the data generated by RNA-SeQC and a custom R-script to produce two metrics, the mtRNA rate and the rRNA rate. Mitochondria reside solely in the cytoplasm, and RNA transcribed from the mitochondrial genome (mtRNA) should not be present in a pure nuclear fraction. The mtRNA rate was defined as the difference in the proportion of reads that mapped to the mtDNA in the nuclear and cytoplasmic fractions of a single sample. This was examined with a view that a positive mtRNA rate would indicate low contamination. To ensure that reads originating from ‘Nuclear mitochondrial DNA sequences’ (NUMTs) were not mistaken for mtRNA mapping reads, we only considered uniquely mapping reads in STAR. The rRNA rate was defined as the difference in the proportion of reads that mapped to ribosomal genes in the nuclear and cytoplasmic fractions of a single sample. As before a positive difference would indicate low contamination within the nuclear fraction, based on the understanding that while rRNA transcription and processing occur in the nucleus (60) assembled rRNAs are transported to the cytoplasm where they contribute to stable ribosome structure.

### ASE signal discovery

ASE signal discovery was performed using a pipeline adapted from Guelfi et al. (50). Briefly, trimmed reads were aligned to personal haploids (created using vcf2diploid (61) (v0.2.6a)) using STAR’s 2-pass alignment in WASP mode with the same parameters as set during the QC alignment described above. The reads that aligned to the parent genomes were then merged using Suspenders tool (62) (v0.2.6) and filtered to select reads that passed the WASP filter. GATK’s (v3.6.0) ASEReadCounter (63) was used to count the reads at the heterozygous sites in the vcf. The vcf was generated by Picard’s (https://broadinstitute.github.io/picard/; Broad Institute) liftover tool. Sites with monoallelic expression were excluded to prevent mis-calling driving ASE signal discovery and only sites showing bi-allelic expression with at least 15 reads were considered valid. The p-value was then calculated using the binomial test, and only hetSNPs with FDR < 5% were considered as a having an ASE signal. The ASE signals were annotated using the Variant Effect Predictor tool (McLaren et al., 2016) (VEP97) with Ensembl cache 97. Variants were assigned to the biotype of the transcript with the most severe consequence. Those with biotypes miRNA, miscRNA, piRNA, rRNA, siRNA, snRNA, snoRNA, tRNA and vaultRNA were grouped under other non-coding RNA (ncRNA) and of the remaining, those not classified as protein-coding, lncRNA or pseudogene were labelled “other”.

In order to identify hetSNPs with significant difference in the allelic ratios between the 2 fractions we also used a statistical approach, applying logit transformation to the allelic ratios and calculating the differential z-scores (64). The p-value was obtained from a 2-tailed test on the z-scores, followed by correction for multiple hypothesis testing. We excluded an outlier hetSNP with highly significant ASE in both fractions and showing highly significant difference in allelic ratios between the fractions. It was located in the gene *RN7SL471P*, with most severe consequence assigned as non coding transcript exon variant by VEP, in the cerebellar cortex tissue of the sample IND001. Functional enrichment analysis was performed using the g:Profiler R package, gprofiler2 (65).

## RESULTS

### Assessment of nuclear and cytoplasmic RNA fractionation quality

RNA fractionation quality was assessed in 22 RNA samples derived from cellular fractions of 3 brain brain regions and originating from four individuals. These samples comprised of nuclear and cytoplasmic RNA fractions of anterior prefrontal cortex (4 individuals), putamen (3 individuals), and cerebellar cortex (4 individuals) (Figure 1a). Firstly, we assessed the expression of genes known to be preferentially localised to the nucleus (*MALAT1*) or cytoplasm (*ACTB*) respectively (66,67). We found an average of 1.6 fold higher expression of *MALAT1* in the nuclear as compared to the cytoplasmic fractions in samples across tissues (Figure 1b) (Wald test p-value, corrected for multiple testing: 1.44E-06 in anterior prefrontal cortex, 2.94E-11 in putamen, 3.70E-04 in cerebellar cortex) and an average of 3.1 fold higher levels of *ACTB* in the cytoplasmic fractions as compared to the nuclear fractions across tissues (Wald test p-value, corrected for multiple testing: 2.03E-22 in anterior prefrontal cortex, 2.26E-20 in putamen, 7.61E-14 in cerebellar cortex). Next, we leveraged reads mapping to rRNA, which could provide a more sensitive measure of fractionation quality. As would be expected, we found that the rRNA rate was consistently higher in the cytoplasmic compared to the nuclear samples with a mean difference of 0.23 in the rRNA rate (paired Wilcox signed rank test p-value = 9.77E-04, Supplementary figure 2a). Finally, we used reads mapping to the mitochondrial genome to assess the quality of RNA fractionation. Since transcription of mtDNA is expected to occur only within mitochondria located in the cytoplasm, reads mapping to the mitochondrial genome but identified within the nuclear RNA-Seq data would indicate cytoplasmic RNA contamination. We found that RNA-Seq data from cytoplasmic fractions had a higher proportion of reads mapping to the mtDNA than their corresponding nuclear samples with a mean difference in the mtRNA mapping of 0.22 (paired Wilcox signed rank test p-value = 9.77E-04, Supplementary figure 2b). We note that we found no evidence of a significant difference in fractionation quality across the three different brain regions analysed (anterior prefrontal cortex, putamen and cerebellar cortex) (Kruskal-Wallis test, p-value = 4.17e-01 for rRNA rate, 4.06e-01 for mtRNA mapping rate).

**Figure 1.**
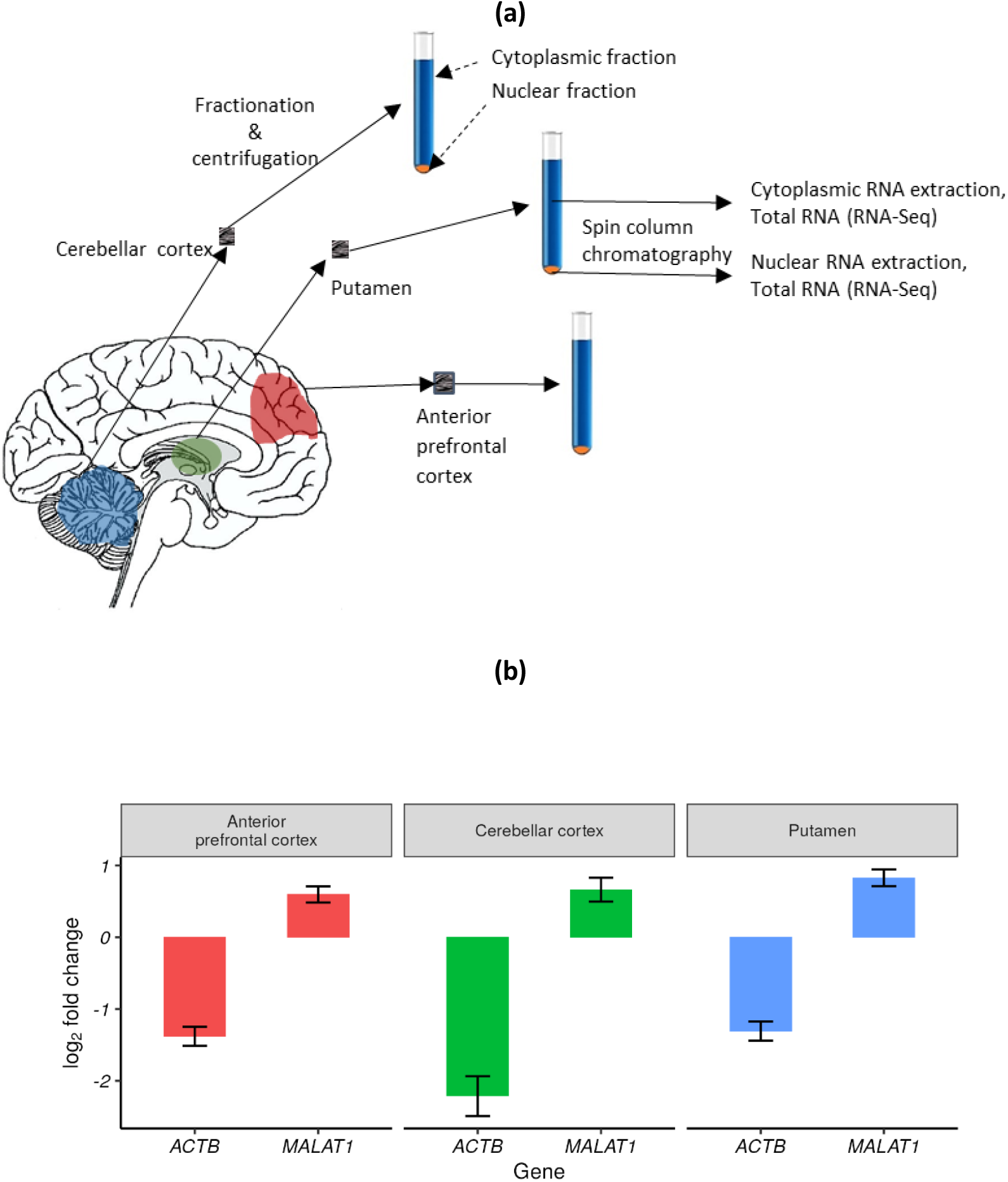
(a) Experimental Overview: The cerebellar cortex (4 individuals), putamen (3 individuals) and anterior prefrontal cortex (4 individuals) regions of the brain, were sampled from human control brains. Separate cellular fractions for each tissue sampled were obtained by centrifugation and RNA extracted using spin column chromatography resulting in a total of 22 nuclear and cytoplasmic samples. (b) Plot showing the quality of fractionation assessed by examining the enrichment of *ACTB* in the cytoplasmic and *MALAT1* in the nuclear fraction. A positive log_2_foldchange indicates gene expression in the nucleus is higher while a negative value indicates the gene expression in the cytoplasm is higher.

### Biotype differences in the transcriptomes derived from nuclear and cytoplasmic RNA

Next, we investigated gene expression within the nuclear and cytoplasmic fractions. We assessed the number of genes detected within each tissue and within each fraction separately. As expected, we found that the vast majority of genes detected in a tissue were present in both fractions and that this was the case for all three tissues (99.9% in anterior prefrontal cortex, 96.96% in putamen, 94.4% in cerebellar cortex). However, we also identified genes which could be detected within all samples in a tissue (based on a normalised count >1), but only in a single fraction (such that the normalised count in the other fraction = 0). Using this approach, we found that the cerebellar cortex had a significantly higher percentage of genes with expression restricted to the nuclear fraction (5.58%, 3-sample test for equality of proportions, p-value = 5.21E-318) as compared to anterior prefrontal cortex (0.09%) and putamen (2.98%). We also observed a very small proportion of genes with expression only in the cytoplasmic fraction (0.004% in anterior prefrontal cortex, 0.06% in putamen and 0.02% in cerebellar cortex). Given that they must be transcribed in the nucleus, this is likely due to expression levels being below detection limits within the nuclear fraction.

Since the majority of genes were robustly detected in both the nuclear and cytoplasmic fractions, we used the paired data to examine differential gene expression in each tissue, controlling for individual level effects. This analysis demonstrated an average of 2,803 genes with significantly higher (FDR <5%) expression in the nucleus and an average of 5,559 genes with significantly (FDR < 5%) higher expression in the cytoplasm (Supplementary file 1). In line with expectation, non-coding RNAs (lncRNAs, pseudogenes) were enriched amongst those genes with significantly higher expression in the nuclear fractions in all tissues (p-value < 2.2E-16). Conversely, protein-coding genes were enriched amongst the genes with significantly higher expression in the cytoplasmic fractions (p-value < 2.2E-16) (Supplementary table 2). Exploring the fold change in expression of the genes which were differentially expressed between the fractions, we observed that as expected, lncRNAs and pseudogenes were differentially expressed with a higher percentage of lncRNA-encoding genes with a fold change in expression of > 2 in the nucleus (in the nucleus vs cytoplasm, 82.5% vs 17.5% in anterior prefrontal cortex, 77.9% vs 22.1% in putamen, 94.9%% vs 5.1% in cerebellar cortex) (Figure 2). These proportions were significantly different between tissues (3-sample test for equality of proportions, p-value = 1.43E-19) with the difference being driven by cerebellar cortex (Benjamani-Hochberg corrected p-values for the two-sample proportion test between anterior prefrontal cortex and cerebellar cortex = 1.43E-06, putamen and cerebellar cortex = 3.67E-17, anterior prefrontal cortex and putamen = 4.57E-01). Conversely, protein-coding genes with >2 fold higher expression within a fraction were primarily located in the cytoplasm (in the nucleus vs cytoplasm, 21.8% vs 78.2% in anterior prefrontal cortex, 24.3% vs 75.7% in putamen, 26.9% vs 73.1% in cerebellar cortex). Again, there was some evidence of a difference in the proportions between tissues (3-sample test for equality of proportions, p-value =4.76E-03). However, the pairwise comparison of proportions showed the significant difference in proportions was between anterior prefrontal cortex and cerebellar cortex (Benjamani-Hochberg corrected p-values for the two-sample proportion test between anterior prefrontal cortex and cerebellar cortex = 1.00E-02, putamen and cerebellar cortex = 1.00E-01, anterior prefrontal cortex and putmen = 2.11E-01). Nonetheless, all tissues had a proportion of protein-coding genes with evidence of >2 fold expression within the nuclear fraction. Given that many protein-coding genes also encode non-coding transcripts this may reflect the retention of such non-coding transcripts within nucleus. Again, this analysis highlighted the cerebellar cortex and the high numbers of lncRNAs differentially expressed within the nuclear fraction.

**Figure 2.**
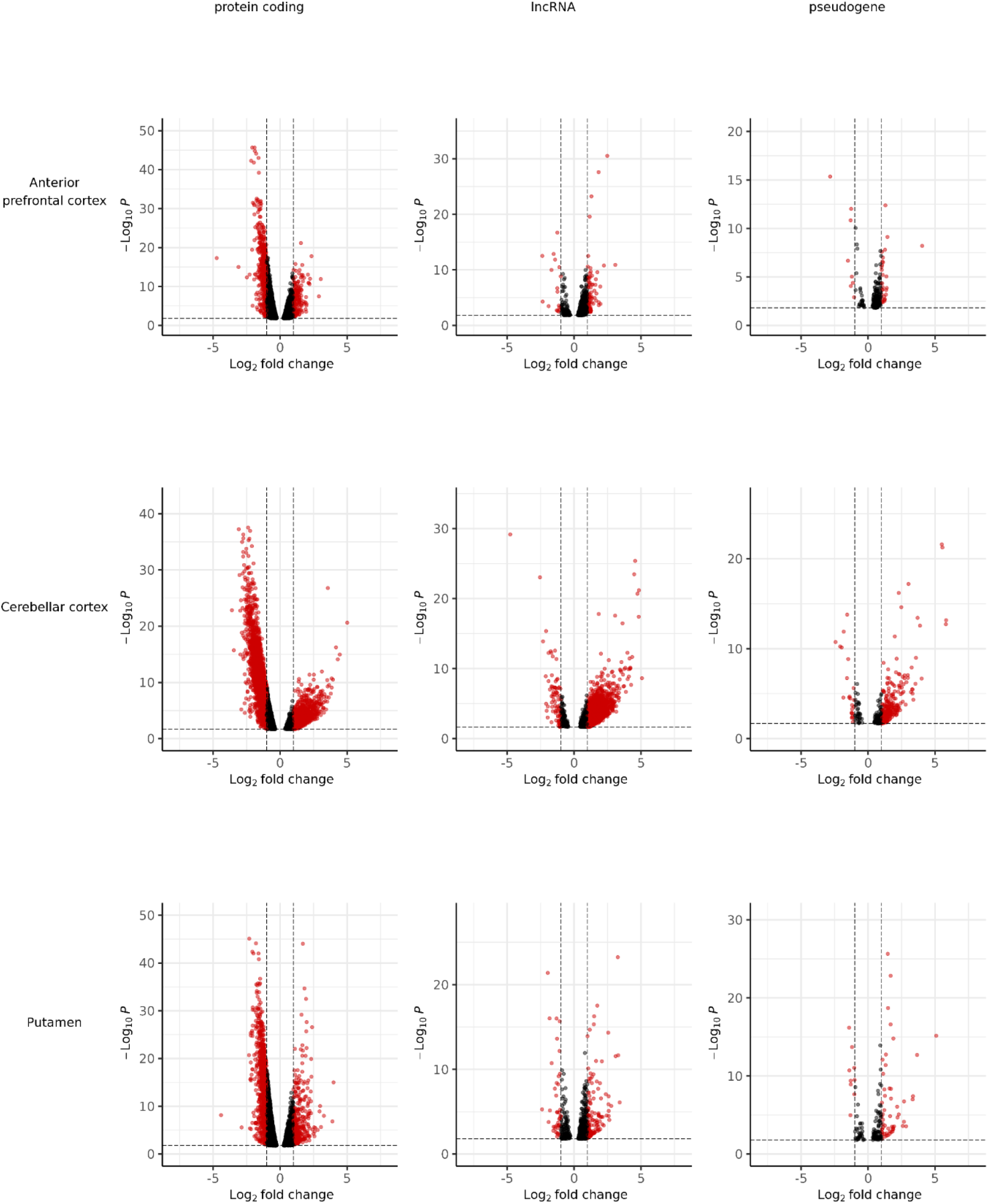
Volcano plots showing the genes with at least double expression (log_2_foldchange >|1|) per tissue. Positive values = expression in the nuclear fraction is higher and negative values = higher expression in the cytoplasmic fraction. 2 outlier datapoints (p-values 7.94E-41 and 5.69E-55) have been excluded from the putamen lncRNA plot. Note : The y-axis scales differ between the plots

Taken together, these findings suggest that nuclear cytoplasmic fractionation enables the capture and analysis of two different transcriptomes that vary significantly in terms of the biotypes of the genes sampled that also extends to the tissue level. In addition, we can see the nuclear transcriptome of cerebellar cortex has an abundance of lncRNAs that is not seen in the other brain tissues.

### Distinct ASE signals across cellular compartments are partially driven by detection and sampling of different transcriptomes

In view of the biotype differences observed in the nuclear and cytoplasmic transcriptomes, we wondered whether there could be related differences in the genetic regulation of gene expression across the cellular compartments. To achieve this, we used ASE analysis which measures differential RNA expression levels between 2 alleles of a heterozygous variant to infer evidence of genetic regulation. As a within individual analysis which naturally pairs the nuclear-cytoplasmic RNA fractionation process within a sample it is ideally suited to this experimental structure. Using this approach, we found that of the 105,341 valid heterozygous SNPs (hetSNPs) studied across all fractions and tissues, an ASE signal was observed in 1.97% at an FDR of < 5% in at least one sample. We observed a significant difference in the number of ASE signals between the tissues (Pearson’s Chi-squared test p-value = 1.47E-39) with the highest percentage of ASE signals observed in cerebellar cortex (1.8%), followed by putamen (1.4%) and anterior prefrontal cortex (0.97%).

The ASE signals observed across all fractions and tissues equated to 1,035 genes representing 9% of all genes studied (and after considering all hetSNPs which could be assigned unequivocally to a single gene). While 261 (25.2%) of the 1,035 genes of interest had a significant ASE signal detected in both fractions, we also identified genes with a significant ASE signal in a single fraction. These ASE-containing genes were termed ‘cytoplasm-specific’ or ‘nucleus-specific’ with the majority being ‘cytoplasm-specific’ (59.8% cytoplasmic, 15% nuclear). Since the distinct ASE signals observed could be due to the specific expression of a gene or transcript within a fraction, we assessed the biotype of the ASE genes identified. We found that of the cytoplasm-specific ASE genes, 4.2% were lncRNAs and 92.9% were protein coding. By comparison, 9.7% of nuclear-specific ASE genes were lncRNAs (2.30-fold higher) and 85.8% were protein coding (Supplementary table 3).

Next, we assessed the effect of SNP location on ASE signal identification. ASE signals were assigned to a genic location, based on their most severe consequence (as defined by Ensembl, Yates et al., 2019). Each hetSNP was assigned the minimum FDR across samples within each fraction and then, as before categorized as ‘both’, ‘cytoplasm-specific’ or ‘nucleus-specific’. While nuclear-specific ASE signals were more commonly located within introns (40.1% nuclear and 12.4% cytoplasmic) (Supplementary table 4), cytoplasm-specific ASE signals were more frequently located within exons (37.7% nuclear and 53.6% cytoplasmic) or untranslated genic regions (UTRs, 22.3% nuclear and 34% cytoplasmic). Overall, these findings were consistent with our expectation that distinct ASE signals within the nuclear and cytoplasmic fractions are driven in part by the differences in the transcriptomes being sampled, with the nuclear fraction being enriched for both lncRNAs and pre-mRNA.

### Distinct ASE signals across cellular compartments are also driven by distinct regulatory processes

Next, we extended our analyses to determine whether some fraction-specific ASE signals were due not only to differences in the RNA content of the two fractions, but also the impact of fraction-specific regulatory processes. With this in mind, the data was examined in a pairwise manner only considering hetSNPs which could be analysed in both fractions of a sample and had a significant signal in at least one of the fractions. Of the 1,852 hetSNPs that could be investigated, 22.7% had ASE signals in both fractions, 63.3% had an ASE signal only in the cytoplasmic fraction and 13.9% had an ASE signal only in the nuclear fraction. Across all tissues, we identified a higher proportion of cytoplasm-specific versus nucleus-specific ASE signals with 5-fold higher cytoplasm-specific as compared to nucleus-specific signals.

Since it is generally assumed that the majority of the genetic regulation of gene expression in human tissues operates through effects on transcription (which occurs in the nucleus), the high percentage of cytoplasm-specific ASE signals was surprising. In order to investigate whether cytoplasm-specific ASE signals were being overestimated due to the application of a p-value cut-off, we re-assessed our data focusing instead on differences in the allelic ratios (defined as the proportion of reference allelic counts) since they are expected to be more robust to the effect of read depth and consequently the power to detect an ASE signal. Using this approach and assigning ASE signals with nuclear-cytoplasmic allelic ratio differences of <=0.1 to the ‘both’ category, we estimated that 22.3% of the hetSNPs of interest were cytoplasm-specific and 7.8% were nucleus-specific (Chi-square goodness of fit p-value= 1.44E-255) (Table 1). This would indicate a 2.84-fold higher percentage of cytoplasm-specific as compared to nucleus-specific ASE signals.

**Table 1.** Percentage of ASE signals (number of signals in brackets), seen in the pairwise analysis, in both and specific fractions. The table includes the percentage of signals after applying the filters for ratio of the total read counts in the nuclear fraction to the cytoplasmic fraction (TRC ratio).

| **ASE signal** | **% ASE signals (*n*)** | **% ASE signals with**<br>**TRC ratio > 0.75 & < 1.25 (*n*)** |
| --- | --- | --- |
| **In both fractions** | 69.9 (1,295) | 60.3 (245) |
| **Cytoplasmic-specific** | 22.3 (412) | 25.9 (105) |
| **Nuclear-specific** | 7.8 (145) | 13.8 (56) |
| **Chi-square goodness of fit**<br>**p-value** | **1.44E-255** | **1.34E-31** |

Given that the use of allelic ratios might be insufficient to account for differences in read depth across the fractions, we further analysed a subset of the 1,852 hetSNPs of interest matching for coverage. More specifically, the only hetSNPs considered were those where the ratio of the total read counts in the nuclear fraction to the cytoplasmic fraction was > 0.75 and < 1.25 (noting that a value of 1 would indicate that the total read count was the same for the hetSNP in both fractions). Following application of these filters, the gene biotypes being studied between the fractions remained unchanged. Despite the stringency of this approach, the number of hetSNPs of interest assigned to the nucleus-specific, cytoplasm-specific and both categories remained significantly different (Chi-square goodness of fit p-value= 1.34E-31) with a higher percentage of cytoplasm-specific ASE signals compared to nucleus-specific signals (25.9% cytoplasm-specific; 13.8% nucleus-specific), and a fold-change of 1.88 (Table 1).

Finally, we used a statistical approach to identify hetSNPs with significant difference in the allelic ratios between the 2 fractions. This was done by application of logit transformation to the allelic ratios and calculation of the differential z-scores (64). Given that only a relatively small number of hetSNPs were tested (N=1,852) we applied a FDR cut-off of 10%. Using this approach and consistent with our previous analyses, we observed a significant difference in allelic ratios in 162 hetSNPs across all tissues, of which the cytoplasm-specific ASEs were still estimated to be close to twice (1.80) as frequent as the nucleus-specific signals. This suggests that a significant amount of common genetic regulation of gene expression is occurring post transcriptionally through cytoplasm-specific processes, such as RNA degradation or localization.

### Compartment-specific ASE signals provide insights into tissue and genic differences in the usage of post-transcriptional genetic regulation of gene expression

Next, we investigated if the identification of distinct regulatory processes in the nuclear and cytoplasmic fractions could provide biological insights. First, we investigated whether ASEs specific to a cellular fraction showed higher tissue specificity, which would indicate tissue differences in the dependence on post-transcriptional genetic regulation of gene expression. To perform this analysis, again we focused on hetSNPs which could be analysed in both fractions of a sample and had a significant signal in at least one of the fractions (1,852 hetSNPs, 868 genes). In order to maximise the number of SNPs analysed, for each tissue and each SNP, we correlated the allelic ratio (defined as the proportion of reference allelic counts) in the two fractions (Figure 3) (Supplementary table 5). Considering ASE signals within the “both” category, we found no significant differences in the correlation between the allelic ratios in each fraction (ANCOVA p-value 0.43). However, in the case of the signals classified as cytoplasm-specific or nucleus-specific there were significant tissue differences in the correlations that were not explained by differences in fractionation quality, though we recognize our sample number is small. This finding was most robust amongst nucleus-specific ASE signals (nucleus-specific ANCOVA p-value 5.67E-04; cytoplasm-specific ANCOVA p-value 2.87E-02) and suggested tissue differences in the overall landscape of genetic regulation of gene expression in brain.

**Figure 3.**
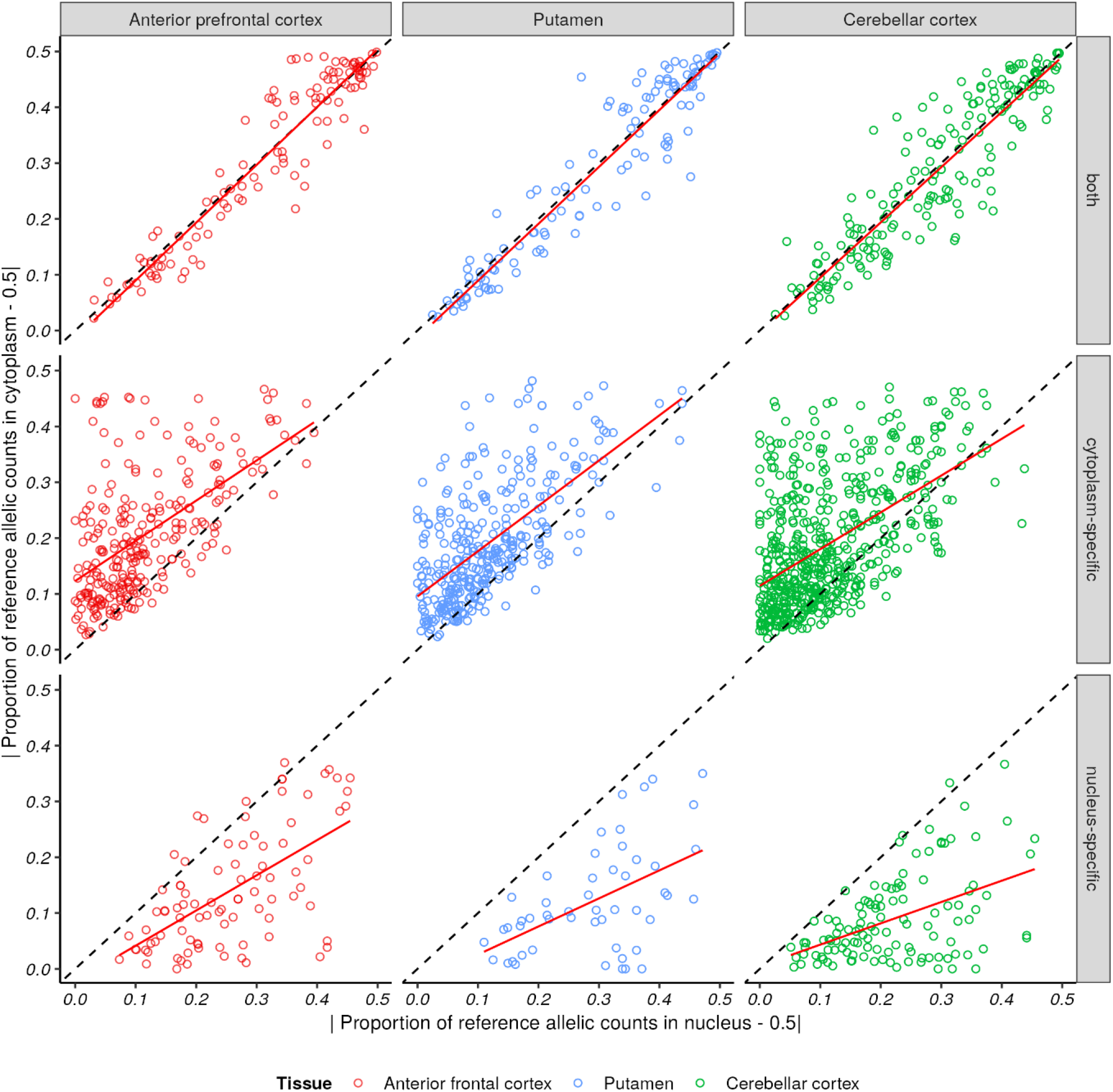
Visualising tissue differences. Plots showing the paired |*allelic ratio* − 0.5| (where allelic ratio is defined as the proportion of reference allelic counts) values for each hetSNP with ASE signals in both fractions, cytoplasm-specific and nucleus-specific. Each point represents a hetSNP in an individual and tissue. The x-axis represents the |*allelic ratio* − 0.5| value for the hetSNP in the nuclear fraction and y-axis its corresponding value in the cytoplasmic fraction. Red line = regression line

Furthermore, we wondered whether genes with evidence of nuclear-or cytoplasm-specific ASE signals differed in their biological functions given the growing evidence for the importance of local regulation of RNA levels in neurons. To address this question, we performed gene ontology enrichment analyses on each gene set combining data across the tissues. We obtained no significant terms for the genes with nuclear-specific ASE signals. However, the genes with cytoplasm-specific ASEs, showed an enrichment of synapse related terms (p-value 0.047) (Supplementary table 6). This finding was driven by genes such as *CAMK2D* and *DLG4*. Interestingly, both genes have been implicated in synaptic plasticity(68-70). While *CAMK2D* is known to be involved in calcium signalling and is crucial for several aspects of plasticity at glutamatergic synapses, *DLG4* encodes a member of the membrane-associated guanylate kinase (MAGUK) family and together with *DLG2* is recruited into postsynaptic NMDA receptor and potassium channel clusters. These findings are highly consistent with the growing literature on the importance of post-transcriptional, local regulation of RNA levels at synapses to enable plasticity(71-73).

## DISCUSSION

Here, we apply ASE analysis to RNA-Seq data derived from paired nuclear and cytoplasmic post-mortem human brain samples to study the genetic regulation of gene expression within each fraction. We demonstrate that in the human brain a significant proportion of genetic regulation of gene expression occurs in the cytoplasm and so post-transcriptionally. We find evidence for tissue and genic differences in usage of post-transcriptional forms of genetic regulation of gene expression, with an enrichment for synaptic genes amongst those with cytoplasm-specific ASE signals. Together, these findings have implications for the interpretation of eQTLs identified through bulk- and single-nucleus RNA-sequencing of human brain, and the use of such eQTL data to identify the biological processes underlying common genetic risk for human brain diseases. In particular, single-nucleus datasets could easily miss any regulatory events that occur in the cytoplasm.

Clearly the robustness of our analyses and resulting conclusions is affected by the quality of RNA extraction from the separate cellular compartments. With this in mind, we used multiple approaches to assess this. In common with the existing literature (67), we began by measuring the expression of genes known to localize specifically to the nucleus (*MALAT1*) or cytoplasm (*ACTB*). As expected, we observed significant enrichments in the appropriate fraction. However, this type of analysis does not sensitively account for the probable leak or contamination of one fraction into the other. Since RNA contamination is expected to occur primarily from the cytoplasm into the nucleus, as the latter only makes up approximately 10-15% of all the cellular RNA (74), we focused on analysing abundant cytoplasmic RNAs, namely rRNA and mtRNA, which are usually considered a form of noise within RNA-Seq datasets. With this in mind, we measured the rRNA rate (defined as reads mapped to rRNA regions/total reads) and mtRNA rate (defined as reads mapped to the mitochondrial genome/total reads) within each compartment, though we recognise that both metrics are likely to overestimate nuclear contamination: rRNA is transcribed within the nucleus, and the presence of transcribed nuclear mitochondrial DNA sequences (NUMTs) could also result in the appearance mtRNA expression within the nuclear fraction. In all samples, we found that the cytoplasmic fractions had a significantly higher rRNA and mtRNA rate compared to the nuclear. Therefore, although there was evidence of contamination, the specific gene enrichments together with the large differences in the rRNA and mtRNA rates between the fractions provided assurance of fractionation quality.

Consistent with expectation, we demonstrated that the transcriptomes derived from each fraction were distinct. Genes detected in only one fraction were primarily a feature of the nuclear transcriptome with the cerebellar cortex having the highest percentage of such genes (5.58% compared to 0.09% in anterior prefrontal cortex and 2.98% in putamen). Although this finding is explained in part by transcription being a nuclear process, it is also likely to be due to the expression and activity of lncRNAs specifically in the nucleus. This conclusion was supported by the fact that differentially expressed lncRNAs were enriched in the nuclear fraction, and that this was most evident in the cerebellar cortex. Thus, this analysis demonstrated that although all tissues had distinct nuclear and cytoplasmic transcriptomes, the nuclear transcriptome of the cerebellar cortex was particularly distinctive. Interestingly, this observation is consistent with a range of reports suggesting a distinct lncRNA expression profile within the cerebellar cortex (75-77).

Given that different transcriptomes were being sampled and analysed, the identification of distinct ASE signals within the two fractions is perhaps unsurprising. Furthermore, it reflected the known enrichment of pre-mRNA and lncRNAs in the nuclear fraction with a higher percentage of intronic ASE signals and lncRNA ASE genes detected in this compartment. However, we noted that the proportion of lncRNA ASE genes within the nuclear fraction was lower than might have been expected given the high numbers of lncRNAs with differential gene expression in the nucleus. While this disparity could suggest a distinct regulatory structure for this transcript biotype, it is most likely to be a function of the technical limitations of sequencing and analysing lncRNAs. This transcript class are expressed at low levels even in the nucleus, and for practical reasons ASE sites were assigned to a transcript and its corresponding biotype based on the most severe consequence of the hetSNP (tending to bias the assignment of SNPs to protein coding transcripts when multiple transcript types are possible).

Since the assessment of ASEs in each fraction separately does not in itself shed light on the regulatory processes unique to each fraction, we also leveraged our paired experimental design to analyse hetSNPs measurable in both fractions of an individual’s tissue. Using this approach, we found that while many of the hetSNPs identified as ASE signals could be detected in both fractions, a significant proportion were only found to be significant in one. Furthermore, most fraction-specific ASE signals were distinct to the cytoplasmic-fraction. This finding remained robust with more stringent forms of analysis, including selecting hetSNPs with very high read depths in both fractions and formally testing the differences in ASEs between fractions using a statistical approach (though the latter was limited to a small number of sites). Importantly, this analysis indicated the existence of different regulatory processes in nuclear versus cytoplasmic fractions as a driver of fraction-specific ASE signals.

Given our current understanding of RNA turnover, ASE signals seen in both fractions could be representative of the impact of genetic variation on transcriptional rate with no additional regulation occurring, while nucleus-specific signals could be indicative of the genetic regulation of RNA transport across the nuclear membrane, and cytoplasm-specific signals an indication of the genetic regulation of a range of post-transcriptional processes, such as RNA stability (78,79) and localization (80-82). Since neurons have extensive projections where specific mRNAs are known to localize and undergo local regulation (83-86), the cytoplasmic ASE signals identified could be generated through incomplete sampling of neurons or synapse-specific processes. While further experimental work is required to investigate these possibilities, the enrichment of cytoplasm-specific ASE genes for synaptic gene ontology terms suggests that these post-transcriptional regulatory processes could be important in human brain tissue and more specifically neurons (73,87). Furthermore, this finding implied that tissue differences in neuronal proportions or types could impact on the balance of nucleus- and cytoplasm-specific ASE signals. Consistent with this hypothesis, the analysis of the allelic ratios identified significant tissue differences in the importance of fraction-specific forms of regulation of gene expression across the tissues sampled (namely anterior prefrontal cortex, putamen and cerebellar cortex). We found that the nucleus-specific and cytoplasm-specific ASE signals were significantly different across the tissues, with cerebellar cortex having weaker correlations of the allelic ratios in both sets of signals.

Taken together, the ASE analysis presented here has implications for our understanding of more commonly available bulk brain eQTL data. While these eQTLs are generally presumed to be regulating transcriptional rate, the significant proportion of cytoplasm-specific ASE signals, which is estimated to be almost double that of the nuclear signals, indicates the importance of post-transcriptional regulation of mRNA through RNA localisation and/or degradation. Although the latter has been analysed using eQTL and ASE-based approaches, to date studies have only been conducted in mice and cell lines (33,88), with this being the first study to specifically explore this question in post-mortem human brain. Similarly, the presence of fraction-specific transcriptomes and ASE signals complicates the interpretation of both single nucleus RNA-sequencing of human brain and related cell type-specific eQTL data, a concern which is not commonly recognised. Thus, we believe the results we have presented both provide novel insights into the regulation of gene expression in the human brain and demonstrate the importance of understanding the molecular processes underlying gene expression regulation.

## DATA AVAILABILITY

Bulk-tissue RNA-sequencing data can be accessed through the European Genome–phenome Archive

## ACKNOWLEDGEMENT

We would like to thank Prof. Michael Simpson (Head of the Genomic Medicine Group at KCL, Head of Human Genetics at Genomics Plc) and his group at KCL for their help in conducting the variant calling on the whole exome sequencing data.

