## Supplemental figures & tables for "Analysis of nucleus and cytoplasm-specific RNA fractions demonstrates that a significant proportion of the genetic regulation of gene expression across the human brain occurs post-transcriptionally"

### Supplementary figures

**Supplementary figure 1.** Principal component analysis plot–Heatmap showing the correlation of the 1st 10 PCs with region, condition, RIN, sex, age and post mortem interval(PMI).

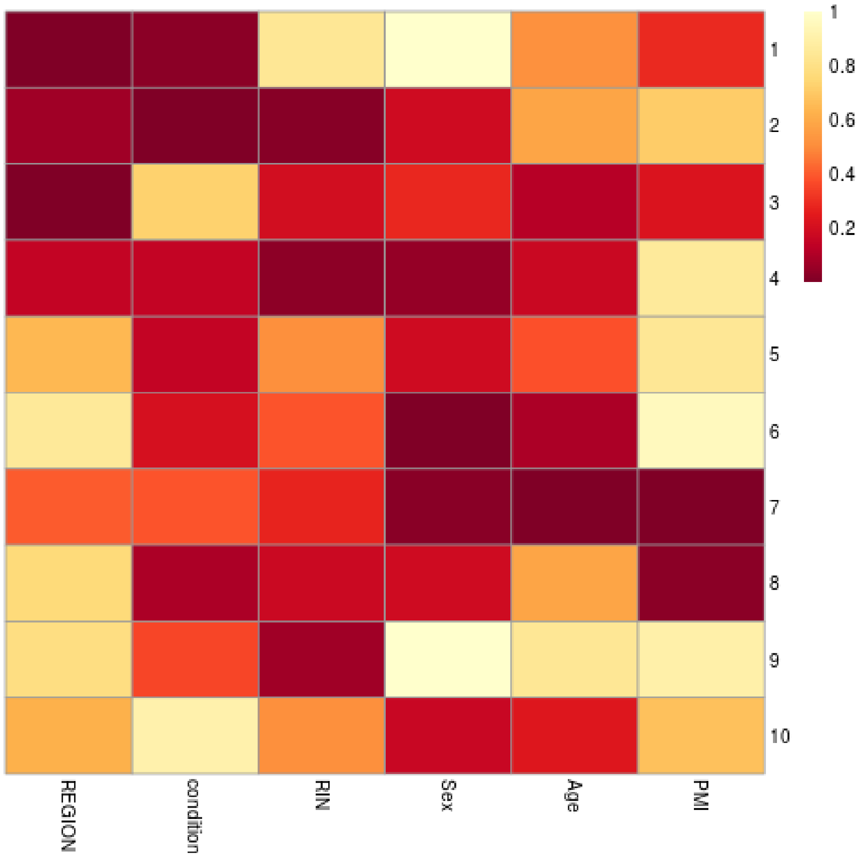

**Supplementary figure 2.** Assessing quality of fractionation by examining the (a) rRNA rate (b) mtRNA mapping rate in the nuclear vs cytoplasmic fractions.

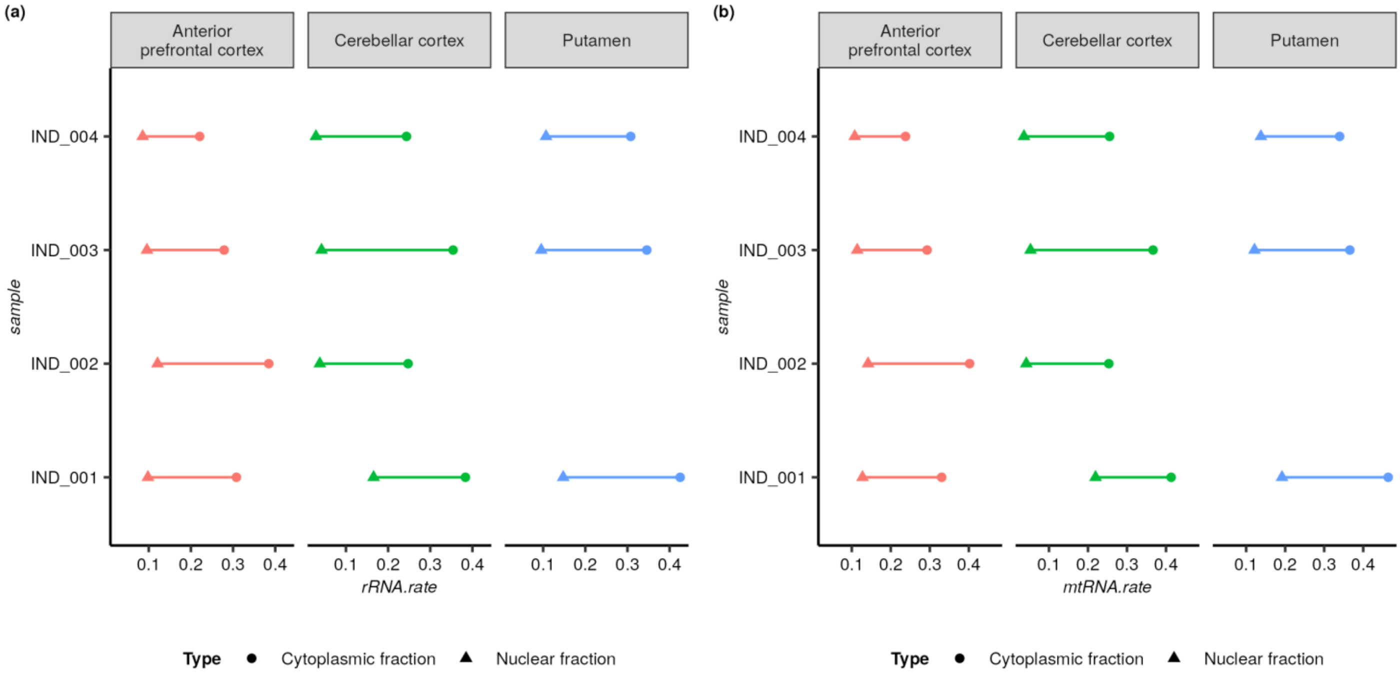

### Supplementary tables

**Supplementary table 1**. Demographic details of individuals

| Individual ID | Sex | Age | Post-mortem interval in hours |
|---|---|---|---|
| IND_001 | Male | 57 | 113 |
| IND_002 | Male | 50 | 41 |
| IND_003 | Male | 53 | 53 |
| IND_004 | Female | 41 | 50 |

**Supplementary table 2.** Percentage of genes with a significant DGE by biotype, in the nucleus and cytoplasm (number of genes are shown within brackets).

Note: The p-values are derived from a 1-sample proportions test with continuity correction as implemented by the prop.test() function in R.

| Tissue | Fraction | Biotypes % | | |
|---|---|---|---|---|
| | | Protein coding | lncRNA | Pseudogene |
| **Anterior prefrontal cortex** | Cytoplasm | 76.8 (4424) | 14.9 (114) | 15.6 (34) |
| | Nucleus | 23.2 (1339) | 85.1 (653) | 84.4 (184) |
| | p-value | < 2.2E-16 | 2.32E-84 | 3.01E-24 |
| **Cerebellar cortex** | Cytoplasm | 80.1 (6310) | 8.4 (197) | 13.4 (58) |
| | Nucleus | 19.9 (1569) | 91.6 (2136) | 86.6 (374) |
| | p-value | < 2.2E-16 | < 2.2E-16 | 3.49E-52 |
| **Putamen** | Cytoplasm | 80.5 (5198) | 22.1 (161) | 22.8 (42) |
| | Nucleus | 19.5 (1256) | 77.9 (567) | 77.2 (142) |
| | p-value | < 2.2E-16 | 3.14E-51 | 1.46E-13 |

**Supplementary table 3.** Distribution of the biotype of genes with an ASE signal in both fractions or fraction-specific (number of genes are shown within brackets).

| Fraction in which ASE signal detected | Biotype (%) | | | |
|---|---|---|---|---|
| | **Protein coding** | **lncRNA** | **Pseudogene** | **Other** |
| Both fractions | 89.3 *(233)* | 5.4 *(14)* | 0.4 *(1)* | 5.0 *(13)* |
| Cytoplasm-specific | 92.9 *(575)* | 4.2 *(26)* | 0.8 *(5)* | 2.1 *(13)* |
| Nucleus-specific | 85.8 *(133)* | 9.7 *(15)* | 0.0 *(0)* | 4.5 *(7)* |

**Supplementary table 4.** Distribution of the most severe consequence of ASE signals by the fraction they were significant in (number of ASE signals are shown within brackets).

| Fraction in which ASE signal detected | Most severe consequence (%) | | |
|---|---|---|---|
| | **Exon variant** | **Intron variant** | **UTR variant** |
| Both fractions | 57.7 *(179)* | 12.9 *(40)* | 29.4 *(91)* |
| Cytoplasm-specific | 53.6 *(490)* | 12.4 *(113)* | 34.0 *(311)* |
| Nucleus-specific | 37.7 *(110)* | 40.1 *(117)* | 22.3 *(65)* |

**Supplementary table 5.** Pearson correlation between the nuclear and cytoplasmic allelic ratios for the ASEs in both fractions, cytoplasm and nuclear specific combining across tissues and by tissue.

| Tissue | Fraction in which ASE signal detected | Pearson correlation |
|---|---|---|
| **Anterior prefrontal cortex** | both fractions | 0.99 |
| | cytoplasm-specific | 0.79 |
| | nuclear-specific | 0.84 |
| **Putamen** | both fractions | 0.99 |
| | cytoplasm-specific | 0.85 |
| | nuclear-specific | 0.80 |
| **Cerebellar cortex** | both fractions | 0.98 |
| | cytoplasm-specific | 0.75 |
| | nuclear-specific | 0.71 |

**Supplementary table 6.** Enrichment analysis - significant terms observed for hetSNPs with a significant difference in allelic ratios and with cytoplasmic-only ASE signals

| Source | Term name | Term size | Term id | p-value | Intersection size | Intersection |
|---|---|---|---|---|---|---|
| REAC | CREB1 phosphorylation through NMDA receptor-mediated activation of RAS signaling | 19 | REAC:R-HSA-442742 | 2.70E-02 | 4 | ENSG00000117676,ENSG00000145349, ENSG00000132535,ENSG00000176884 |
| REAC | Ras activation upon Ca2+ influx through NMDA receptor | 15 | REAC:R-HSA-442982 | 4.74E-02 | 3 | ENSG00000145349,ENSG00000132535, ENSG00000176884 |
| REAC | Negative regulation of NMDA receptor-mediated neuronal transmission | 14 | REAC:R-HSA-9617324 | 4.74E-02 | 3 | ENSG00000145349,ENSG00000132535, ENSG00000176884 |
| REAC | Unblocking of NMDA receptors, glutamate binding and activation | 16 | REAC:R-HSA-438066 | 4.74E-02 | 3 | ENSG00000145349,ENSG00000132535, ENSG00000176884 |
| REAC | Long-term potentiation | 16 | REAC:R-HSA-9620244 | 4.74E-02 | 3 | ENSG00000145349,ENSG00000132535, ENSG00000176884 |
| REAC | Neurotransmitter receptors and postsynaptic signal transmission | 114 | REAC:R-HSA-112314 | 4.74E-02 | 8 | ENSG00000145863,ENSG00000117676, ENSG00000176533, ENSG00000109158, ENSG00000145349,ENSG00000132535, ENSG00000176884,ENSG00000162989 |
